# Exploring glioma heterogeneity through omics networks: from gene network discovery to causal insights and patient stratification

**DOI:** 10.1101/2024.07.02.601715

**Authors:** Nina Kastendiek, Roberta Coletti, Thilo Gross, Marta B. Lopes

## Abstract

Gliomas are primary malignant brain tumors with a typically poor prognosis, exhibiting significant heterogeneity across different cancer types. Each glioma type possesses distinct molecular characteristics determining patient prognosis and therapeutic options. This study aims to explore the molecular complexity of gliomas at the transcriptome level, employing a comprehensive approach grounded in network discovery. The graphical lasso method was used to estimate a gene co-expression network for each glioma type from a transcriptomics dataset. Causality was subsequently inferred from correlation networks by estimating the Jacobian matrix. The networks were then analyzed for gene importance using centrality measures and modularity detection, leading to the selection of genes that might play an important role in the disease. Spectral clustering based on patient similarity networks was applied to stratify patients into groups with similar molecular characteristics and to assess whether the resulting clusters align with the diagnosed glioma type. The results presented highlight the ability of the proposed methodology to uncover relevant genes associated with glioma intertumoral heterogeneity. Further investigation might encompass biological validation of the putative biomarkers disclosed.

## 1. Introduction

Gliomas are primary malignant brain tumors with a typically bad prognosis, accounting for 80% of malignancies in the brain [1]. Due to their large heterogeneity, gliomas encompass different cancer types, where each type possesses distinct characteristics that influence patient prognosis and therapeutic options. The World Health Organization (WHO) Classification of Tumors of the Central Nervous System (CNS) has been changing throughout the years. In the most recent version of the WHO CNS tumor classification [2], the glioma types are classified mainly based on the sample’s molecular profiles instead of histological features. Gliomas in adults are characterized by three types: astrocytoma, oligodendroglioma, and glioblastoma [2]. The latter exhibits a wildtype status for the IDH genes, and it represents the most common and aggressive type, with a median survival of only 14 months [3]. Conversely, oligodendrogliomas and astrocytomas present mutations in at least one IDH gene, being molecularly distinguished by the presence/absence of the combined loss of the short arm chromosome 1 and the long arm of chromosome 19 (1p/19q codeletion), respectively [2].

Studying gliomas on a molecular level is essential for understanding the biological mechanisms behind these brain tumors, ultimately advancing cancer medicine. Omics data comprise various layers of biological information, e.g. genomics (DNA sequences), transcriptomics (RNA transcripts), proteomics (protein abundances), methylomics (DNA methylation level), and metabolomics (metabolite profiles). This type of data is increasingly available with technological advances, e.g., through high-throughput RNA sequencing and protein mass spectrometry.

Network science is highly recognized in the field of cancer research to study the complexity of diseases and extract relevant biological information [4]. Biological networks help to explain the disease by studying the interactions between biological entities such as genes. In gene co-expression networks, the nodes represent genes that are connected with an edge if the corresponding genes are significantly co-expressed in the RNA sequencing (RNA-seq) data across different samples. Analyzing these networks can uncover key genes that have strong connections to other genes in the network, as well as hub genes that strongly interact with several others. Another type of network in the study of cancer is the patient similarity network, in which patients are represented as nodes and the similarities between them as edges connecting the nodes. These networks are used for clustering patients into groups with similar molecular characteristics, which may be indicative of common underlying biological mechanisms. As patients may respond differently to treatments, patient stratification is critical for precision medicine approaches, ensuring that patients with similar characteristics receive the most effective treatments, adapted to their specific type of tumor.

In a previous study, Martins et al. [5] explored the potentiality of applying network discovery and clustering techniques to glioma data. Glioma patients were grouped according to 2016-WHO classification guidelines, and undirected networks were used to perform variable selection before applying the K-means clustering. The results pointed out some discordance between clusters and 2016-WHO classes, fostering further studies based on the new 2021-WHO diagnostic label assignments.

This study presents a comprehensive network-based approach to glioma patient data to investigate their complex gene expression profiles, leading to the identification of potentially important genes and patient stratification. For this, RNA-seq glioma data from The Cancer Genomics Atlas (TCGA) were used, with patients grouped according to the updated 2021-WHO glioma types [6]. From these vast datasets, the variables (i.e., genes) were preselected using the corresponding TCGA proteomics dataset containing a set of proteins in the major biochemical pathways in cancer, i.e., the genes encoding for the proteins present in the proteomics dataset were retained for further analysis. A gene co-expression network for each tumor type was constructed using the graphical lasso method to reveal the interplay of genes in each glioma type. Next, the obtained network structure was used to infer causality among genes through the Jacobian matrix estimation, a crucial step providing insights into the direction of gene interactions. Centrality measures and modularity detection were then applied to the constructed networks to identify central genes as potential biomarkers of each glioma type. As a final step, spectral clustering was applied to patient similarity networks based on the selected variables from the gene networks to stratify patients into distinct groups and evaluate whether these patient groups matched the clinically diagnosed glioma types.

## 2. Materials and Methods

### 2.1 Gene co-expression networks via graphical lasso

The graphical lasso is a statistical method for estimating the graphical structure in a Gaussian graphical model. It is particularly useful in high-dimensional scenarios where the number of variables is higher than the number of samples, as it induces sparsity in the model. The objective is to find a sparse graph that represents the conditional independence structure among the variables by applying a lasso penalty to the inverse covariance matrix. Suppose n multivariate normal observations of dimension p, and let **∑** and **S** be the empirical and theoretical covariance matrix, respectively. The graphical lasso, as implemented by [7], applies a lasso penalty to the inverse covariance matrix **Θ** = **∑**^-1^ by solving the optimization problem

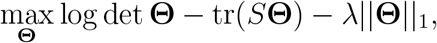

where λ is the regularization parameter and ||**Θ**||_1_ is the L1 norm of **∑**^-1^ (sum of the absolute values of all its elements).

If element ij in the estimated inverse covariance matrix **∑**^-1^ is zero, then variables i and j are conditionally independent [7]. The non-zero elements quantify the sign and strength of the direct relationship between the corresponding pair of variables, while controlling for the influence of all other variables. The inverse covariance matrix thus provides information about the partial correlation structure of the nodes [8].

The regularization term is the penalty that induces sparsity in the model. The sparsity reflects the assumption that not all variables are directly related to each other, and pushes entries in the inverse covariance matrix to zero. A larger value of λ enforces more sparsity. Here, the Stability Approach to Regularization Selection (StARS) is used for selecting the regularization parameter λ. StARS assesses the variation in network topology across different subsamples of the data and optimizes λ to use the minimum necessary regularization to ensure the network’s reproducibility under random sampling [9].

### 2.2 Causal discovery by Jacobian reconstruction

Inferring causality from observed correlations, i.e., moving from undirected to directed networks, is a key challenge. In omics studies, it holds the promise to unveil relevant links driving tumorigenesis. When a system is subject to some noise, it is possible to employ the theory of stochastic processes showing that correlations emerging from samples can be interpreted as a ‘signature’ of the underlying deterministic system [10]. This leads to a relationship between the covariance of the samples and the system’s Jacobian matrix, which is derived according to [10] and [11] as follows:

Given a system of N variables defined by ***X*** = (*X*_1_, …, *X*_*N*_)^T^, their response to small fluctuations around an equilibrium can be approximated as

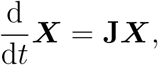

with the Jacobian **J** of dimension *N*. The system with noise can be modelled by the Langevin-type equation

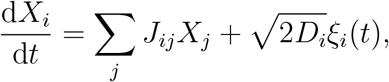

where *ξ*_*i*_(*t*) is Gaussian white noise, with zero mean and unit variance and *D*_*i*_ is the mean amplitude of the fluctuations of *X*_*i*_. By using the corresponding stationary Fokker-Planck equation for the probability distribution, the Lyapunov equation is obtained [12]:

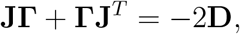

where **Γ** is the covariance matrix with entries r_*ij*_ = ⟨*X*_*i*_*X*_*j*_⟩ and **D** is the fluctuation matrix, describing the internal noise of the system.

Based on this systematic relationship, it is possible to recover the deterministic system by inferring the system’s Jacobian matrix from the sample data, if additional knowledge about the system is available [10].

Note that the Jacobian matrix is generally non-symmetric, whereas the covariance matrix is always symmetric. This leads to a number of constraints in the Lyapunov equation that is lower than the degrees of freedom of the Jacobian, making it impossible to determine causality from correlation alone. However, if additional information, such as the topology of interactions, is available, the full Jacobian can be reconstructed. Barter and co-authors [11] propose a step-by-step method to solve the Lyapunov equation for **J**, by considering additional constraints. Specifically, it is needed to set *J*_*ij*_ = 0 for *N*(*N*™1)/2 pairs of nodes *i* and *j*, forcing the absence of the corresponding variable relations [11].

### 2.3 Biomarker selection through node centrality and modularity detection

Centrality measures in network analysis are used to identify the most important nodes within a network. There are various measures that each offer different perspectives on a node’s importance based on its connections and position within the network structure. Here, the strength and eigenvector centrality are used, which are both calculated from a non-negative adjacency matrix **A** describing relations among a set of *N* nodes with centralities *c*_1_, …, *c*_*N*_.

The strength of a node is quantified by summing the weights of the edges the node i is connected to [13]

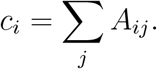

With strength centrality, it is presumed that a node with significant or several interactions has a higher level of importance in the network.

In the eigenvector centrality, a node’s centrality is determined by summing its connections to others, whereby each connection is weighted by the other node’s centrality

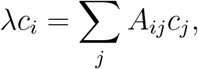

where λ is a constant required so that the equations have a non-zero solution.

This equation can be expressed in matrix notation

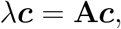

where λ and ***c*** represent, respectively, an eigenvalue and the associated eigenvector of the matrix **A**. Among the possible eigenvalues of **A**, the leading one (i.e., the eigenvalue with the largest real part) is considered, since its eigenvector is guaranteed to have positive entries (Perron–Frobenius theorem) [14]. Then, the *N* components of the leading eigenvector represent the centralities of each node. In this concept of centrality, the importance of a node is proportional to the importance of its neighboring nodes, creating a recursive relationship where connections to highly central nodes enhance a node’s own centrality [15].

In directed networks, it must be distinguished between outward and inward centrality. Outward centrality quantifies the influence of a given node over the network, while inward centrality reflects how much a node is influenced by the others [16]. With an adjacency matrix where rows reflect the out-degrees and columns the in-degrees of nodes, outward strength centrality is calculated by summing over the rows and inward strength centrality is computed by summing over columns. For eigenvector centrality, outward centrality calculates the right eigenvector of **A**, satisfying *λ****c*** = **A*c*** while inward centrality calculates the left eigenvector, satisfying *λ****c***^*t*^ = ***c***^*t*^**A** [16]. Consequently, outward centrality can be calculated similarly to undirected networks and inward centrality is calculated based on the transpose of **A**. For this study, outward centrality was calculated, following the notion that genes that exert a high influence over the network could be potentially important drivers in the disease process.

In modularity detection, the task is to organize a network into modules, or communities, by grouping nodes that have a similar role in the network. This can be regarded as a way to select groups of functionally related genes instead of focusing only on single nodes. Stochastic Block Models (SBMs) provide a probabilistic approach for this purpose [17].

A SBM is a generative model that is based on a structure of *N* nodes divided into *B* groups. The partition is given by a vector ***b***, where each entry *b*_*i*_ ∈{1, …, *B*}defines the group membership of node i. Given the partition of the nodes ***b***, the idea is generating a network in which the nodes are grouped accordingly, by maximizingthe probability *P*(**A|*b***), where **A** is the adjacency matrix. The probability *P*(**A**|***b***) determines the likelihood of edges existing between nodes based on their group memberships. If Bayesian inference approach is used, we can revert the problem, and infer the most probable modularity structure that could have generated an observed network, by computing *P*(***b*|A**) through the Bayesian rule [18]. The number of groups B and the most likely partition ***b*** can be computed in combination by optimizing the Integrated Classification Likelihood [19].

SBMs lead to relatively homogeneous degree distributions within modules, which might not always align with networks estimated from real-world data. To address this limitation, Degree-corrected SBMs (DcSBMs) account for variations in node degrees within the same module [18].

### 2.4 Spectral clustering of patient similarity networks

Spectral clustering is a powerful method for grouping complex data into distinct clusters by partitioning a similarity graph. It is particularly useful for clustering samples based on high-dimensional omics data. In a patient similarity network, the samples to be clustered are regarded as nodes of a graph. The edges between the nodes represent the similarities between the samples, which are typically evaluated using the Euclidean distance across all variables. Taking the nearest neighbor graph to capture the local structure of the data is especially important for detecting non-linear structures in the data.

Spectral clustering captures the manifold structure of complex data structures through the eigendecomposition of the graph’s Laplacian matrix. The eigenvectors corresponding to the smallest non-zero eigenvalues form a new, lower-dimensional, feature space where the data points are separated into distinct clusters [20]. John and co-authors [21] proposed the *Spectrum* method for spectral clustering of complex omics data, implemented as an R package. It constructs the patient similarity matrix from the omics data, computes the nearest neighbor graph, reduces noise to better reveal underlying structures, and finally performs the spectral clustering. For more details of the method, refer to [21].

### 2.5 Data and analysis workflow

The transcriptomics and proteomics data were retrieved from GDAC Broad Firehose (https://gdac.broadinstitute.org), a portal collecting data generated by The Cancer Genome Atlas (TCGA) [22]. In particular, we considered normalized RNA-seq and protein abundance data from the TCGA-GBM and TCGA-LGG projects [23, 24, 25]. These datasets group the glioma patients into glioblastoma and lower-grade glioma (LGG) in line with the 2007 WHO classification [26]. LGGs aggregate astrocytoma, oligodendroglioma, and oligoastrocytoma samples. The dataset was updated to align with the 2021 WHO classification, thereby reallocating oligoastrocytoma cases to one of the three defined glioma types [6].

The transcriptomics data, derived from RNA sequencing, provides the gene expression levels of over 20,000 genes. The proteomics data, obtained through reverse phase protein arrays (RPPA), is a functional proteomics dataset accounting for nearly 200 proteins involved in major biochemical signaling pathways in cancer [27]. For the computational feasibility of the methods employed in this study, a dimensionality reduction of the transcriptomics dataset was necessary. Therefore, the analysis was specifically focused on the expression levels of those genes from the transcriptomics dataset that encode the proteins contained in the proteomics dataset, according to the map provided by the GDAC Broad Firehose portal. This resulted in a significant variable reduction while still ensuring that important genes involved in regulating cancer pathways were included in the analysis. This subset of the transcriptomics dataset (transcriptomics_*S*_) comprises 145 RNA-seq variables measured over 206 samples for glioblastoma, 255 samples for astrocytoma, and 166 samples for oligodendroglioma. This reduced dataset was normalized by a high-dimensional Gaussian copula with nonparametric marginals (non-paranormal), which transforms the variable distribution to achieve normality [28]. The nonparanormal normalization was performed by the *huge*.*npn* R function. Only the transformed variables normally distributed in all glioma types according to the Jarque-Bera test (*jarque*.*test* function from the *moments* R package) were used in the network inference process.

A preliminary spectral clustering using the *Spectrum* method was performed to get insights into the biological information contained in the reduced transcriptomics_*S*_ dataset compared to the full transcriptomics dataset and the proteomics dataset. The *Spectrum* R tool was employed using its default settings, with the cluster number set to *c* = 3 to match the three glioma types, i.e., glioblastoma, astrocytoma, and oligodendroglioma.

A first network discovery step is part of the methodological workflow to infer association and causal networks for the three glioma types, using the graphical lasso method and the Jacobian matrix estimation, respectively. Figure 1 describes the analysis workflow. The undirected networks were inferred from the transcriptomics data separately for each glioma type by the graphical lasso method using the *huge* R function. The StARS method was employed using the *huge*.*select* R function to determine the optimal regularization parameter *λ* for each of the three networks. The resulting values for *λ* were closely aligned across all types, with *λ* ≃ 0.3, leading to a similar level of sparsity in the graphs. Based on the inverse covariance matrices, undirected gene co-expression networks were constructed for each glioma type, providing information about the partial correlation structure of the gene relations at the transcriptomics level. Owing to the induced sparsity, some genes had no connections, being excluded for further analysis.

**Figure 1:**
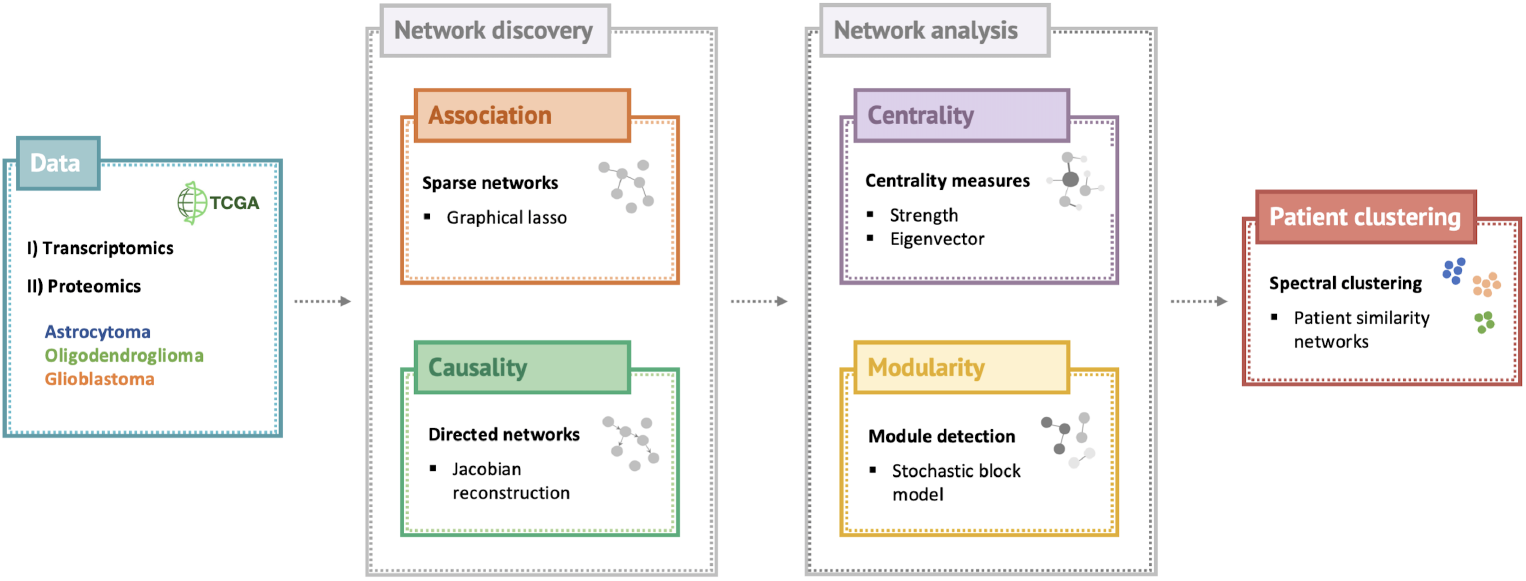
Schematic representation of the analysis workflow encompassing network discovery, network analysis, and patient clustering applied to TCGA glioma transcriptomic and proteomics datasets.

To infer causality among gene relations, the Jacobian matrix was computed for each glioma type, following the step-by-step method according to [11]. The fluctuation matrix describing the internal noise of the system is assumed diagonal with 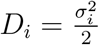, for *i* = 1,…, *N*, where 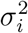 are the variances of the individual variables [12]. The structural information needed as additional constraints was retrieved from the sparsity of the gene co-expression networks previously estimated. The zero entries in the inverse covariance matrix 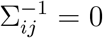 defined the zero entries in the Jacobian matrix J_*ij*_ = 0. For each glioma type, a directed network was deduced based on the respective Jacobian matrix, describing how one gene influences another. Self-loops and the sign of the edge weights were ignored for further analysis since the identification of important genes requires focusing only on the strength of the relationship between the genes. All graphs shown were visualized using the force-directed layout algorithm by Fruchterman and Reingold, which positions nodes that are directly connected closer to each other and increases the distance of isolated nodes [29].

A second step of the analysis workflow was the analysis of the networks inferred, based on centrality measures and modularity detection. The strength and eigenvector centrality were calculated to determine the importance of the genes in the glioma networks, while the modularity structure was computed to reveal functionally related groups of genes that may be important. For modularity detection via DcSBMs, the *greed* R function was used. If the adjacency matrix of the network is supplied, the function computes the most likely modularity structure obtained by DcSBM [19].

In the proposed methodology, relevant genes were selected from the undirected and directed networks, by identifying the genes covering the top 10% of importance in the networks and the most important gene from each module, based on strength and eigenvector centrality. This leads to eight sets of potential biomarkers for each glioma type.

In a final step of the analysis, spectral clustering using the *Spectrum* method was applied to the resulting glioma transcriptomics datasets accounting for the genes selected by the network analysis step, to test whether these reduced datasets allow the distinction of known glioma types. The Adjusted Rand Index (ARI) [30] was computed to quantify how much the identified clusters were in agreement with the actual updated labels. The quality of the clustering in terms of tightness and separation was evaluated by the Average Silhouette Width (ASW) [31]. In the *Spectrum* method the first c = 3 eigenvectors were used for clustering the data; thus, the feature space based on those was used to calculate the ASW.

## 3. Results

To assess if the biological information of the considered omics data is capable of capturing glioma intertumoral heterogeneity, a preliminary unsupervised study was performed. In particular, the *Spectrum* clustering method was applied to the full transcriptomics dataset (16,217 features), the proteomics dataset (174 features), and the transcriptomics_*S*_ dataset (143 features), i.e., covering only genes coding for the proteins collected in the proteomics dataset.

The confusion matrices of the clustering results together with the two clustering performance measures ASW and ARI are provided in Table 1. The full RNA transcriptomics dataset produced the best clustering outcome, with the highest scores in both measures, suggesting these data reflect glioma heterogeneity. In contrast, the proteomics dataset lead to poor-quality clusters, not matching the known glioma types. This outcome indicates that the proteomics dataset used was not informative for glioma patient stratification. Interestingly, the clustering obtained for the proteomics-informed dataset, i.e., transcriptomics_*S*_, achieved performances comparable to those obtained from the full transcriptomics dataset. Moreover, the lower input dimension massively reduced the computational complexity, making this subset of features a valuable input dataset for our analysis.

**Table 1:** Results from spectral clustering based on the transcriptomics and proteomics datasets (transcriptomics_*S*_ stands for the proteomics-informed transcriptomics dataset)

|  | Transcriptomics<br>16217 features |  |  | Proteomics<br>174 features |  |  | Transcriptomics<sub>S</sub><br>143 features |  |  |
| --- | --- | --- | --- | --- | --- | --- | --- | --- | --- |
| Astrocytoma | 4 | 239 | 12 | 57 | 42 | 107 | 4 | 243 | 8 |
| Glioblastoma | 190 | 15 | 1 | 33 | 14 | 69 | 181 | 24 | 1 |
| Oligodendroglioma | 0 | 6 | 160 | 37 | 49 | 57 | 0 | 17 | 149 |
| ARI | 0.82 |  |  | 0.01 |  |  | 0.75 |  |  |
| ASW | 0.71 |  |  | 0.45 |  |  | 0.65 |  |  |

Figure 2 shows the resulting patient similarity network derived from the transcriptomics_*S*_ dataset. The graph shows a remarkable distinction between the three glioma types, despite some samples being closer to others from a different class. For instance, few glioblastoma cases are allocated in the area of astrocytoma and oligodendroglioma, suggesting some molecular affinities that might deserve further investigation. Overall, astrocytoma seems to share more similarities with the other two classes, while oligodendroglioma and glioblastoma can be better distinguished. This prior result is in line with biological knowledge since astrocytoma is characterized by the same cell type as glioblastoma [32], while having mutations in the same gene family [2].

**Figure 2:**
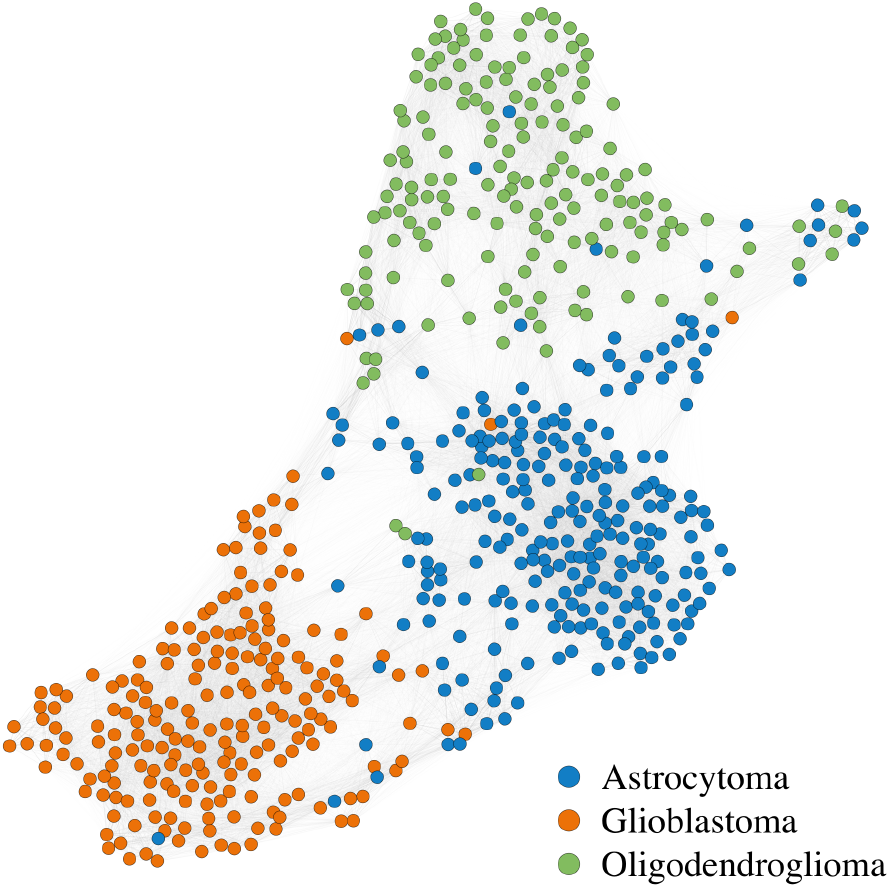
Patient similarity network computed from the proteomics-informed transcriptomics dataset transcriptomics_*S*_. Each node represents a patient, and colors are assigned based on the associated glioma type. The nodes are placed using the force-directed layout algorithm by Fruchterman and Reingold [29].

### 3.1 Network Discovery and Analysis

The estimation of the undirected glioma networks by the graphical lasso method was performed separately for glioblastoma, astrocytoma, and oligodendroglioma. The resulting gene co-expression networks comprised a set of genes that have at least one connection, i.e., 141 genes for glioblastoma, 143 genes for astrocytoma, and 141 genes for oligodendroglioma. Based on these network structures, the Jacobian matrices were computed for each type. We recall that the Jacobian matrix is not symmetric, as two genes can influence each other in both directions and in different ways (*J*_*mn*_*≠J*_*nm*_). Moreover, the entries can be positive or negative, where a positive entry *J*_*mn*_ means that gene *m* has an activating influence on gene *n* while a negative entry indicates an inhibiting influence. However, the sign of the influence was not taken into account in our analysis; instead, our focus was on the direction and strength of the relationships between the genes which is reflected in the directed networks.

Given the difficulty of a visual analysis of these dense networks, centrality measures and module detection were used to detect key genes as potential biomarkers. To this aim, for each glioma type, both undirected and directed networks were considered. The modularity detection algorithm applied for the undirected networks of astrocytoma, oligodendroglioma and glioblastoma types identified 8, 7, and 5 modules, respectively. In the directed networks, 4 modules were detected for both astrocytoma and oligodendroglioma, and 9 modules for glioblastoma.

The detection of such modules and the computation of gene importance improves the interpretability of the overall networks. As an example, Figures 3 and 4 show the astrocytoma undirected and directed network, respectively, where the identified modules are highlighted with different colours. In both figures, each node size is proportional to the gene importance, based on strength (subfigures a) and eigenvector (subfigures b) centralities.

**Figure 3:**
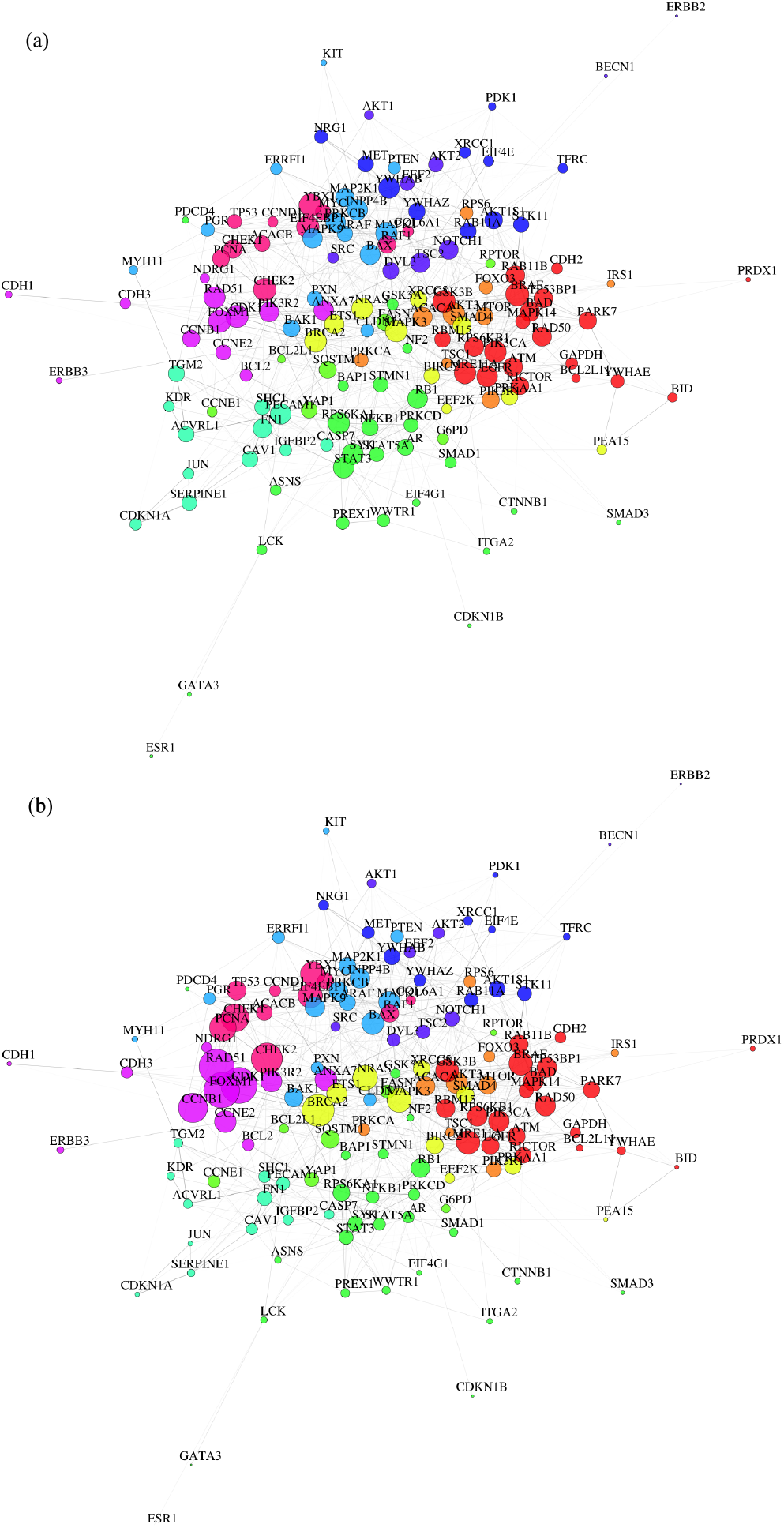
Astrocytoma undirected networks with node color based on the modularity structure, and node size proportional to their importance within the networks based on (a) strength centrality and (b) eigenvector centrality.

**Figure 4:**
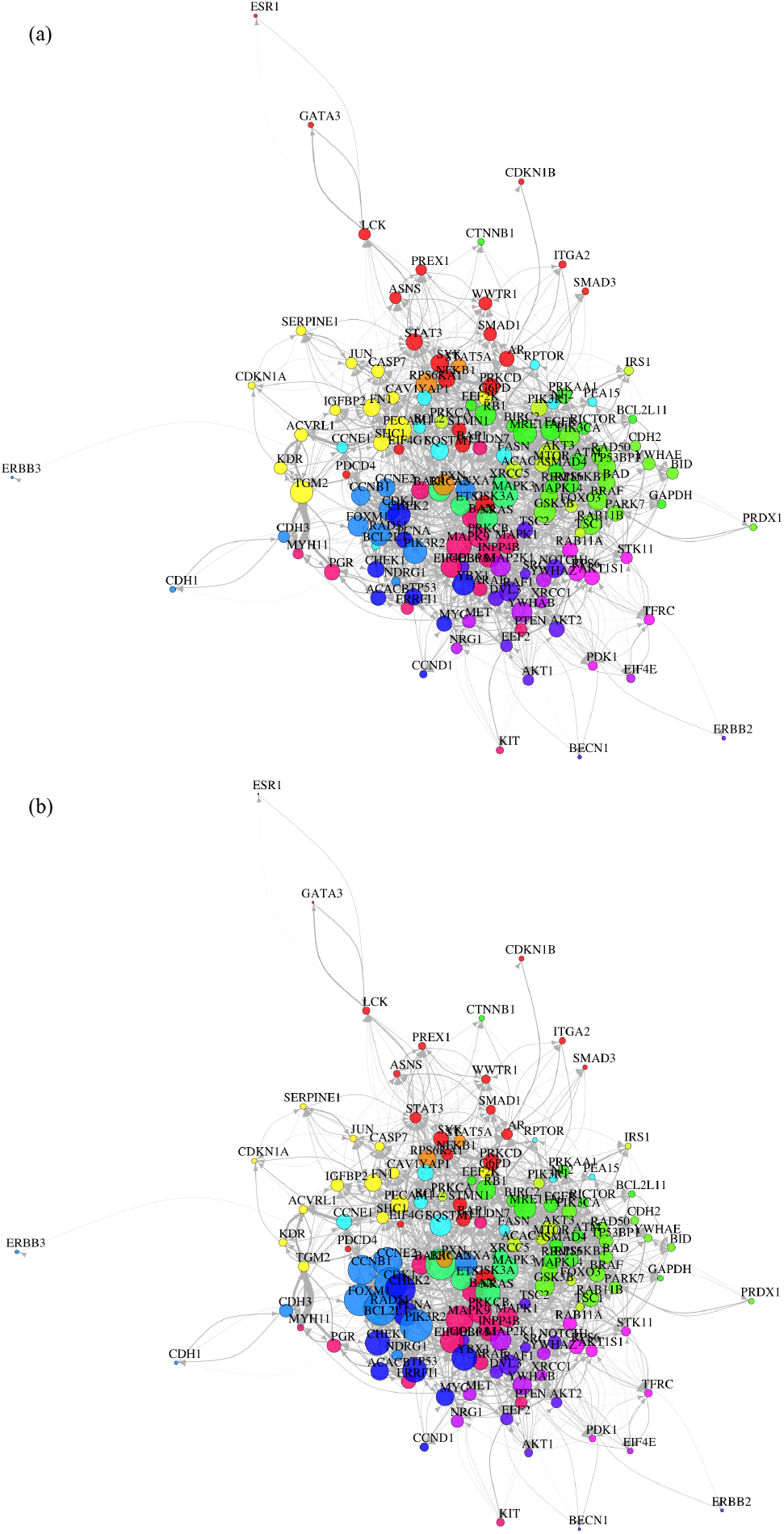
Astrocytoma directed networks with node color based on the modularity structure, and node size proportional to their importance within the networks based on (a) strength centrality and (b) eigenvector centrality.

From the observation of Figures 3 and 4 emerges that eigenvector centrality assigns a more distinct ranking among the genes, with some nodes significantly larger than others, whereas strength centrality returns a more uniform value to each gene, resulting in many nodes of similar size. As a consequence, coupling module detection with eigenvector centrality offers a more interpretable outcome, as it allows the detection of entire groups of genes with high importance within the network. For instance, it is possible to detect the most relevant module based on eigenvector centrality in both undirected and directed astrocytoma networks, i.e., the purple module in Figure 3b and the blue module in Figure 4b, respectively. Figure 5 focuses on those subnetworks, allowing a deeper discussion of the included gene relations. Interestingly, despite a generally different module structure of directed and undirected networks, the most important module in each of these two cases is mostly constituted by the same genes, with the only exception of *BCL2*, which is exclusively present in the purple module of the undirected astrocytoma network. This high node overlap among the two graph representations allows the disclosure of how the estimated gene relationships change from the undirected to the directed network. We observe that the edge weights of the undirected module appear relatively uniform, while varying significantly in the directed network. In this case, many pairs of nodes have links in both directions, yet their weights highly differ, suggesting a dominant influence of one gene over the other. For instance, from Figure 5b, *PIK3R2* is predicted to have a considerable impact on *RAD51*, influencing it by a direct strong positive link, and by a path mediated by *FOXM1*, which might indicate an important gene regulatory process.

**Figure 5:**
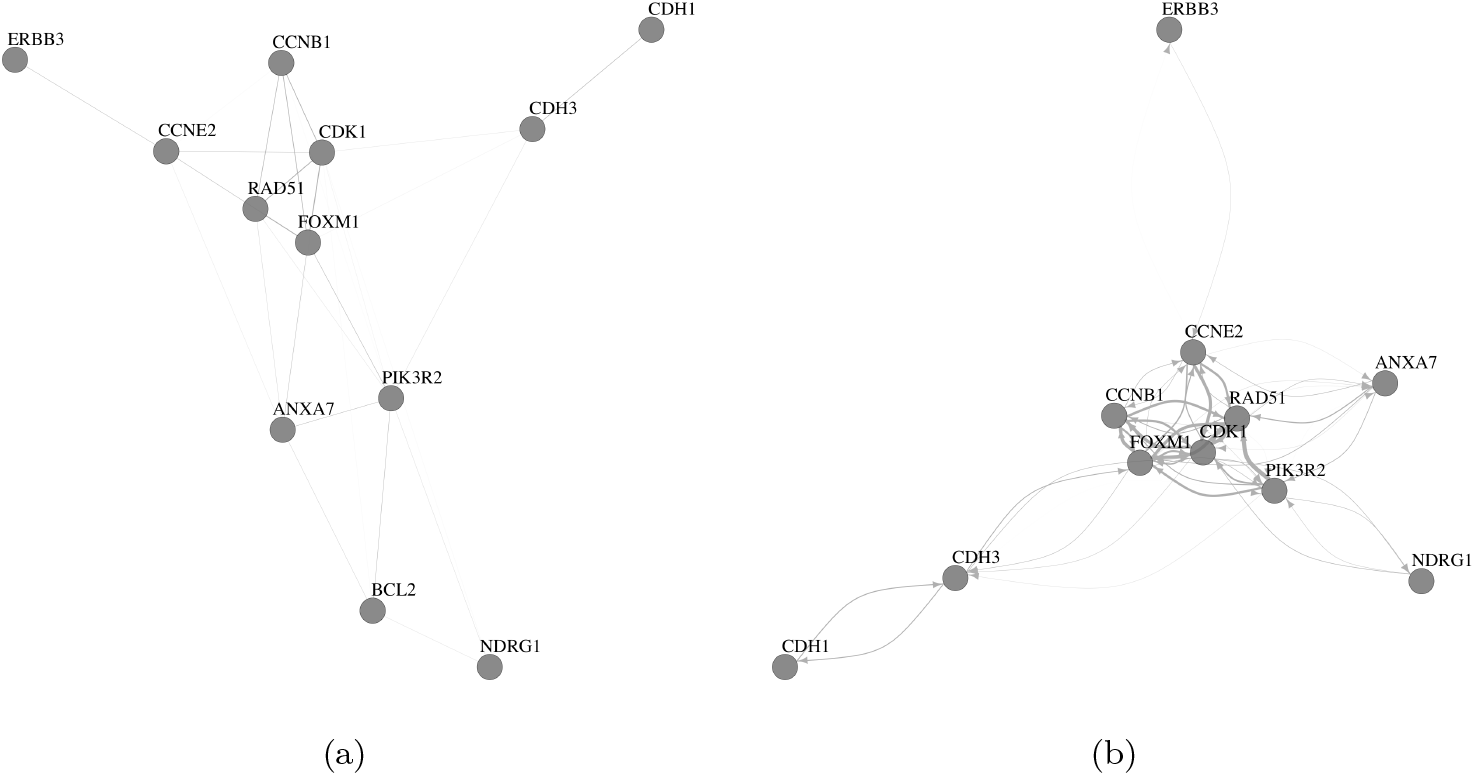
Most relevant module in astrocytoma (a) undirected network and (b) directed network. These modules include the same genes, with the only exception of *BCL2*, which is exclusively present in the module of the undirected astrocytoma network. The nodes are placed using the force-directed layout algorithm by Fruchterman and Reingold [29].

A literature review on these genes revealed findings aligning with our results, highlighting a potential common ground in DNA repair mechanisms. Specifically, *PIK3R2* regulates the activity of the enzyme Phosphoinositide 3-kinase (PI3K), which is linked to many cell functions, such as growth, proliferation, and cell motility. This gene is widely known in cancer, where it is frequently mutated, and associated with tumor proliferation and increased invasion [33, 34]. Due to these characteristics, recent studies proposed *PIK3R2* as a cancer prognostic marker [35, 36].

On the other hand, *FOXM1* is a transcription factor gene involved in cell proliferation, self-renewal, and tumorigenesis [37]. Commonly upregulated in cancer [38], it is considered a prognostic marker and a therapeutic target [39].

The existence of a link going from *PIK3R2* to *FOXM1* is supported by literature, as *FOXM1* activation is controlled by PI3K/ATK pathway [40], which is crucial in glioma progression [41].

Instead, *RAD51* has its primary function in DNA-damage repairs [42], a process extremely important in cancer, and broadly investigated in glioma due to its implication in treatment failure. In particular, it has been shown that higher levels of *RAD51* are associated with poor glioma survival, due to its role in repairing DNA damages caused by radiation and chemotherapy, contributing to treatment resistance and tumor recurrence [43, 44].

Many studies pointed out the *FOXM1* role in regulating DNA damage response [45, 46, 47]. In pediatric glioma, it has been shown that inhibition of PI3K reduces the DNA repair functions by a pathway involving *FOXM1*. In particular, *FOXM1* is responsible for activating promoters of genes having DNA damage repair functions, such as *RAD51* [48]. In ovarian cancer, the depletion of *PIK3R2* results in increased DNA damage and a reduction of the proteins encoded by the *RAD51* gene [49].

All these findings suggest that the genes identified in our approach deserve further exploration to understand their biological functions in glioma. If such regulations are confirmed by biological validations, we may postulate that the *PIK3R2* gene might impact glioma therapy resistance by alternating DNA repair processes, with important clinical implications.

For potential biomarker detection and further biological interpretation, the following sets of genes were selected from the networks: (set 1) top 10% of importance in undirected networks based on the strength centrality, (set 2) top 10% of importance in undirected networks based on the eigenvector centrality, (set 3) most important per module in undirected networks based on strength centrality, (set 4) most important per module in undirected network based on eigenvector centrality, (set 5) top 10% of importance in the directed networks based on strength centrality, (set 6) top 10% of importance in directed networks based on eigenvector centrality, (set 7) most important per module in the directed networks based on strength centrality, (set 8) most important per module in directed networks based on eigenvector centrality. The selected genes for each glioma type from all eight sets are listed in Table 2.

**Table 2:**
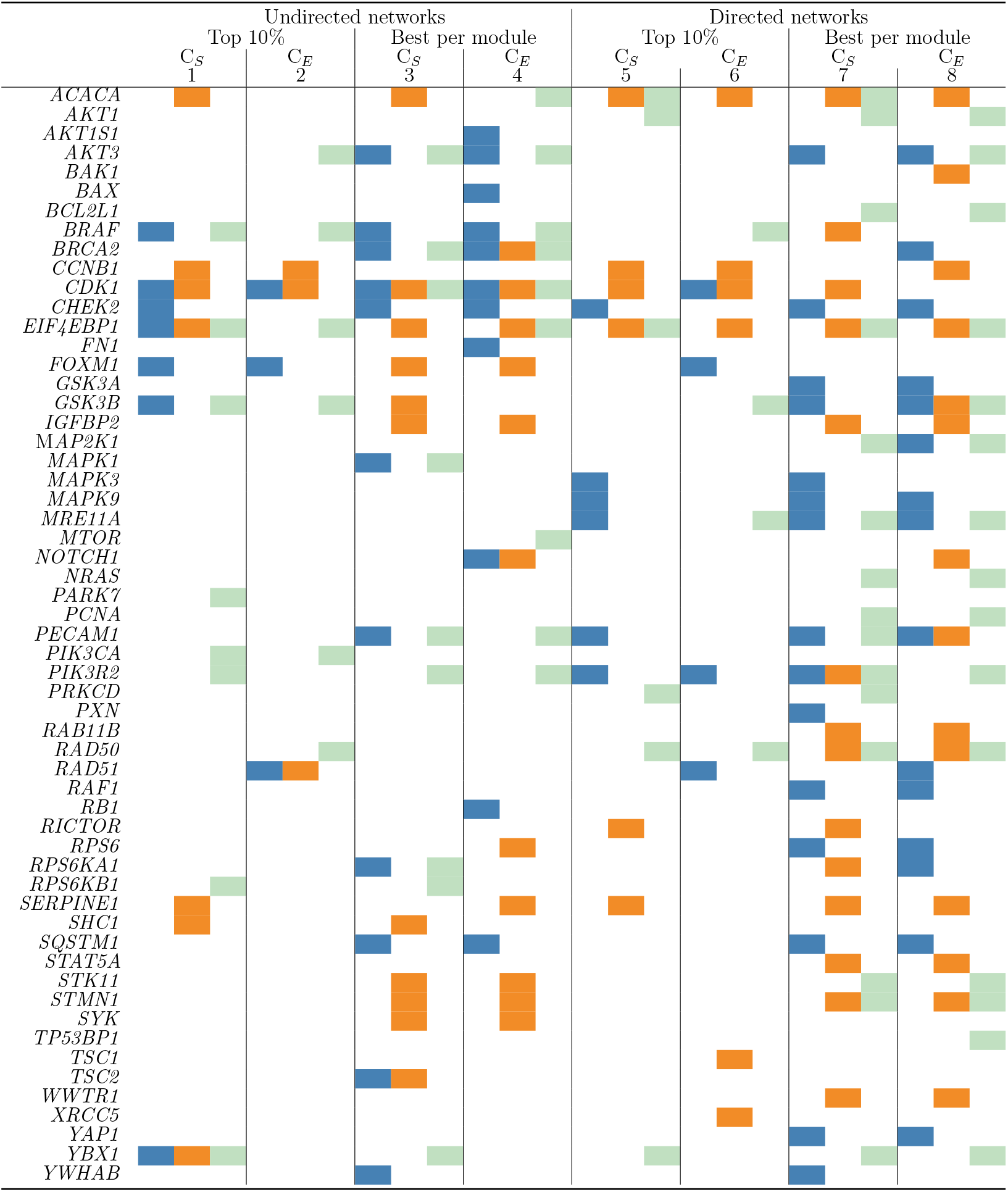
Selected genes in the 8 sets: undirected networks vs. directed networks, genes covering the top 10% of importance vs. most important gene per module, strength centrality (C_*S*_) vs. eigenvector centrality (C_*E*_)(Astrocytoma, Glioblastoma, Oligo-dendroglioma)

The overview provided in Table 2 allows the identification of promising genes that might be biomarker candidates of glioma heterogeneity. Indeed, many of them are detected as key variables for a single glioma type. For instance, *CHEK2* is exclusively detected for astrocytoma from the directed and undirected network in 6 out of 8 sets, suggesting an important role in the disease. Analogously, *CCNB1* is only selected for glioblastoma, while *RAD50* is consistently selected for oligodendroglioma from the directed network estimation. On the other hand, there are genes detected as important for two or even all the glioma types, such as *CDK1, EIF4EBP1* and *PIK3R2*, which might indicate their relevance in processes shared across different types, and therefore being also suitable as potential markers.

### 3.2 Patient Clustering

Since the dataset exploration performed as a first stage of our analysis assessed that the initial dataset was suitable to perform spectral clustering, we would like to preserve this ability after variable selection. Therefore, as a final step, clustering was performed by considering only data from the selected features. The confusion matrices of the clustering results together with ASW and ARI values are given in Table 3.

**Table 3:** Results from spectral clustering based on the variables selected from undirected vs. directed networks; genes covering the top 10% of importance vs. most important gene per module; strength centrality (C_*S*_) vs. eigenvector centrality (C_*E*_))

| Undirected networks |  |  |  |  |  |  |  |  |  |  |  |  |
| --- | --- | --- | --- | --- | --- | --- | --- | --- | --- | --- | --- | --- |
|  | Top 10% |  |  |  |  |  | Best per module |  |  |  |  |  |
| | $C_S$ | | | $C_E$ | | | $C_S$ | | | $C_E$ | | |
|  | 15 features<br>(set 1) |  |  | 10 features<br>(set 2) |  |  | 23 features<br>(set 3) |  |  | 23 features<br>(set 4) |  |  |
| Astrocytoma | 24 | 50 | 181 | 21 | 171 | 63 | 26 | 204 | 25 | 31 | 207 | 17 |
| Glioblastoma | 170 | 4 | 32 | 140 | 45 | 21 | 186 | 16 | 4 | 194 | 12 | 0 |
| Oligodendroglioma | 4 | 145 | 17 | 4 | 135 | 27 | 4 | 46 | 116 | 9 | 12 | 145 |
| ARI | 0.47 |  |  | 0.20 |  |  | 0.51 |  |  | 0.64 |  |  |
| ASW | 0.55 |  |  | 0.42 |  |  | 0.51 |  |  | 0.63 |  |  |

| Directed networks |  |  |  |  |  |  |  |  |  |  |  |  |
| --- | --- | --- | --- | --- | --- | --- | --- | --- | --- | --- | --- | --- |
|  | Top 10% |  |  |  |  |  | Best per module |  |  |  |  |  |
| | $C_S$ | | | $C_E$ | | | $C_S$ | | | $C_E$ | | |
|  | 16 features<br>(set 5) |  |  | 13 features<br>(set 6) |  |  | 36 features<br>(set 7) |  |  | 35 features<br>(set 8) |  |  |
| Astrocytoma | 18 | 235 | 2 | 46 | 124 | 85 | 7 | 246 | 2 | 11 | 217 | 27 |
| Glioblastoma | 139 | 66 | 1 | 186 | 5 | 15 | 165 | 40 | 1 | 179 | 26 | 1 |
| Oligodendroglioma | 2 | 70 | 94 | 11 | 119 | 36 | 1 | 65 | 100 | 1 | 141 | 24 |
| ARI | 0.36 |  |  | 0.30 |  |  | 0.51 |  |  | 0.38 |  |  |
| ASW | 0.59 |  |  | 0.43 |  |  | 0.52 |  |  | 0.54 |  |  |

Overall, the clustering performance decreased compared to the one provided by the initial dataset. However, despite the considerable variable reduction, most of the final subsets lead to high-quality clusters, with ASW score higher than 0.5. Also, the ability of class recovery was good for half of the subsets, with ARI values around 0.5 or higher. The highest ARI and ASW values were obtained by considering the most important genes from the modules in the undirected graphs according to eigenvector centrality (set 4), suggesting this subset of genes is the most promising to capture glioma heterogeneity.

Generally, the clustering performance was improved when considering the modularity structure of the networks. However, this result might be also affected by the number of genes included in the analysis, which is generally larger than the one deriving from the top 10% approach. The worst performances in terms of ARI and ASW values were provided by considering genes from the top 10% based on eigenvector centrality. Nevertheless, if the number of genes included in the analysis might affect clustering results, it is worth to note that both these cases are characterized by the smallest number of features.

The confusion matrices reveal that oligodendroglioma and astrocytoma are often assigned to the same cluster. This outcome aligns well with the known similarities in the genetic profiles of these glioma types. Astrocytomas and oligodendrogliomas are both low-grade gliomas exhibiting mutations in the same gene family, i.e., the IDH genes. Glioblastomas, on the other hand, display a wild-type IDH status, and are characterized by genetic or histological alterations, uncommon to astrocytomas and oligodendrogliomas. In diagnosis, this IDH mutation is a crucial marker that distinguishes astrocytomas and oligodendrogliomas, from the more aggressive glioblastomas [2]. This outcome is also confirmed by previous studies based on the same datasets, investigating glioma heterogeneity by network inference, which also highlights the similarity between astrocytoma and oligodendroglioma [5, 50]. Furthermore, there is more overlap between glioblastoma and astrocytoma than between glioblastoma and oligodendroglioma, which is reasonable since both glioblastoma and astrocytoma arise from the same cell type, namely astrocytes.

The identified genes could play significant roles in the glioma disease due to their importance in the networks. In this study, clustering was used as an unsupervised approach to assess the effectiveness of the selected genes in differentiating the glioma classes. An additional literature review on the subset of genes leading to the best clustering results (set 4) was conducted to explore their specific functions in gliomas and cancer in general. The findings can be found in Table A.1. In the literature review, all selected features are mentioned in relation to human cancers, some in the context of gliomas. Among studies about gliomas, many studies are focused on glioblastoma, widely studied due to being the most common and aggressive type of glioma.

## 4. Conclusions

In this study, we propose a network-based methodology to discover relevant gene interaction networks associated with different glioma types. The methodology encompasses estimating gene co-expression networks from transcriptomics data using the graphical lasso method, estimating causal relationships by the Jacobian matrix, and the analysis of gene importance based on centrality measures and modularity detection. Further clustering of patient similarity networks evaluates the suitability of selected relevant network genes to stratify the patients into the diagnosed glioma types.

While the sparse gene co-expression networks identified by the graphical lasso method stand as valuable information regarding the molecular associations involved in different glioma types, the causal links disclosed through the Jacobian matrix further increase the biological understanding of the directions of the corresponding interactions. Although these cannot be considered as direct gene regulatory networks due to the absence of transcription factors that usually mediate gene-gene interplay [51], the estimated causal networks provide an important indication of gene interactions, a crucial starting point to foster further research.

To solve for the Jacobian matrix, the sparsity in the edges obtained via the graphical lasso estimation was used as constraints. Further research might consider other prior information available from curated databases regarding known molecular interactions involved in glioma or cancer in general.

The analysis of the resulting networks through node centrality and modularity detection provided valuable insights into gene importance and the structure of gene interactions in the glioma networks and led to a selection of genes as potential biomarkers. The ability of these genes to group patients into the known glioma types was assessed by spectral clustering of patient similarity networks. Overall, a general agreement in the resulting cluster composition and the established glioma types was achieved.

A set of promising genes was identified in gliomas from a mathematical and computational standpoint. We conducted a literature review to evaluate previous reports on the role of the selected genes in glioma or cancer in general, which is essential to assess their potential for therapy research. In particular, by analyzing the directed gene relations in a astrocytoma module, we were able to detect very promising regulatory mechanisms, which aligns with prior studies, yet not comprehensively investigated in glioma. The confirmation of these findings might increase the overall glioma understanding, with potentially important clinical implications. Genes not yet associated with glioma might potentially be considered for experimental validation, which involves biologically testing the most promising candidates to examine their role in the development, progression, and therapy of glioma.

## Acknowledgements

This work was supported by national funds through Fundação para a Ciência e a Tecnologia (FCT) with references CEECINST/00042/2021, UIDB/00297/ 2020 and UIDP/00297/2020 (NOVA Math, Center for Mathematics and Applications), UIDB/00667/2020 and UIDP/00667/2020 (UNIDEMI), and the research project PTDC/CCI-BIO/4180/2020 “MONET – Multi-omic networks in gliomas” (doi: 10.54499/PTDC/CCI-BIO/4180/2020). The results presented here are based upon data generated by the TCGA Research Network: https://www.cancer.gov/tcga.

## Appendix A. Biomarker candidates

**Table A.1:** Selected genes and their function in gliomas and cancer in general.

| Gene | Function |
| --- | --- |
| *ACACA* | Aberrant expression increases cancer risk [52]; down-regulation suppresses malignant progression of prostate cancer [53] |
| *AKT1S1* | Regulates the PI3K/AKT/PRAS40/mTOR pathway; involved in several cancer cell processes [54, 55] |
| *AKT3* | Amplification promotes glioma progression [56]; it delays tumor progression, therefore strategies that inhibit *AKT3* may be unhelpful in some glioblastoma patients [57] |
| *BAX* | Important regulator in controlling cell proliferation and apoptosis [58]; over-expression of *BAX* is a target gene therapy of glioma [59] |
| *BRAF* | Lower-grade gliomas of childhood are characterized by *BRAF* mutations [60]; *BRAF* mutation associated with improved survival in glioma [61] |
| *BRCA2* | Downregulation can sensitize glioma cells to killing by anti-cancer drugs [62]; factor for poor prognosis in glioma patients [63] |
| *CDK1* | Inhibiting CDK1 activity promotes apoptosis in glioblastoma [64]; significantly enriched in glioblastoma cell cycle [65] |
| *CHEK2* | Activated upon DNA damage; Associated with breast cancer [66]; *CHEK2* gene polymorphism might correlate with prognosis of glioblastoma patients [67] |
| *EIF4EBP1* | Overexpression linked to poor survival and disease progression in hepatocellular carcinoma [68]; increased expression in malignant gliomas [69] |
| *FN1* | Upregulated in Glioblastoma; it promotes Glioblastoma cell proliferation by altering *PTPRM* methylation [70] |
| *FOXM1* | Promotes glioma cells progression [71]; overexpressed in malignant glioma [72] |
| *IGFBP2* | *IGFBP2* expression predicts IDH-mutant glioma patient survival [73]; it promotes glioma tumor stem cell expansion and survival [74] |
| *MTOR* | Therapeutic target for glioblastoma [75]; important role in glioblastoma [76] |
| *NOTCH1* | *NOTCH1* expression is upregulated in glioma and associated with tumor progression [77]; prognostic factor for patients with glioma [78] |
| *PECAM1* | Downregulated in Glioblastoma; correlated to the overexpression of HIF-1α, a transcriptional regulator increasing tumor aggressiveness, invasiveness and resistance to radiotherapy and chemotherapy [79] |
| *PIK3R2* | It regulates the activity of PI3K, which is linked to many cell functions, and well known in glioma [80, 81, 82]; frequently mutated, and associated with tumor proliferation and increased invasion [33, 34]; proposed as a cancer prognostic marker [35, 36] |
| *RB1* | Downregulated in human brain tumors [83]; alteration of the RB1-pathway related genes are associated with shorter survival in low-grade gliomas [84]; in Glioblastoma, loss of RB1 expression correlates with hypermethylation, which is not reported in low-grade astrocytoma [85] |
| *RPS6* | Upregulated in high-grade glioma; involved in several mechanisms characterizing malignant cancer behaviour [86, 87, 88] |
| *SERPINE1* | Identified as regulator of glioblastoma cell dispersal [89]; biomarker for diffuse lower-grade gliomas [90] |
| *SQSTM1* | Acts as a regulator in autophagy in Glioblastoma cells [91], and cancer in general [92] |
| *STK11* | Potential tumour suppressor gene [93]; mutations occur particularly in lung and colorectal cancer [94] |
| *STMN1* | Overexpression associated with the prognosis of meningioma patients [95]; overexpressed in many types of human malignancies [96] |
| *SYK* | SYK inhibition blocks proliferation and migration of glioma cells [97]; associated with malignant phenotype and immune checkpoints in diffuse glioma [98] |

## References

[1] M. Weller, W. Wick, K. Aldape, M. Brada, M. Berger, S. M. Pfister, R. Nishikawa, M. Rosenthal, P. Y. Wen, R. Stupp, G. Reifenberger, Glioma, Nature Reviews Disease Primers 1 (1) (2015) 15017. doi:10.1038/nrdp.2015.17.

[2] D. N. Louis, A. Perry, P. Wesseling, D. J. Brat, I. A. Cree, D. Figarella-Branger, C. Hawkins, H. K. Ng, S. M. Pfister, G. Reifenberger, R. Soffietti, A. von Deimling, D. W. Ellison, The 2021 WHO classification of tumors of the central nervous system: a summary, Neuro-Oncology 23 (2021) 1231–1251. doi:10.1007/s00401-016-1545-1.

[3] S. Mohammed, D. M, A. T, Survival and quality of life analysis in glioblastoma multiforme with adjuvant chemoradiotherapy: a retrospective study, Reports of Practical Oncology and Radiotherapy 27 (6) (2022) 1026–1036. doi:10.5603/RPOR.a2022.0113.

[4] M. B. Lopes, E. P. Martins, S. Vinga, B. M. Costa, The role of network science in glioblastoma, Cancers 13 (5) (2021) 1045. doi:10.3390/cancers13051045.

[5] S. Martins, R. Coletti, M. B. Lopes, Disclosing transcriptomics network-based signatures of glioma heterogeneity using sparse methods, BioData Mining 16 (1) (2023) 26. doi:10.1186/s13040-023-00341-1.

[6] M. L. Mendonça, R. Coletti, C. S. Gonçalves, E. P. Martins, B. M. Costa, S. Vinga, M. B. Lopes, Updating tcga glioma classification through integration of molecular profiling data following the 2016 and 2021 who guidelines, bioRxiv (2023). doi:10.1101/2023.02.19.529134.

[7] J. Friedman, T. Hastie, R. Tibshirani, Sparse inverse covariance estimation with the graphical lasso, Biostatistics 9 (3) (2008) 432–441. doi:10.1093/biostatistics/kxm045.

[8] J. S. Hawe, F. J. Theis, M. Heinig, Inferring interaction networks from multi-omics data, Frontiers in Genetics 10 (2019) 535. doi:10.3389/fgene.2019.00535.

[9] H. Liu, K. Roeder, L. Wasserman, Stability approach to regularization selection (stars) for high dimensional graphical models (2010). doi:10.48550/ARXIV.1006.3316.

[10] R. Steuer, J. Kurths, O. Fiehn, W. Weckwerth, Observing and interpreting correlations in metabolomicnetworks, Bioinformatics 19 (8) (2003) 1019–1026. doi:10.1093/bioinformatics/btg120.

[11] E. Barter, A. Brechtel, B. Drossel, T. Gross, A closed form for jacobian reconstruction from time series and its application as an early warning signal in network dynamics, Proceedings of the Royal Society A: Mathematical, Physical and Engineering Sciences 477 (2247) (2021) rspa.2020.0742, 20200742. doi:10.1098/rspa.2020.0742.

[12] N. Van Kampen, Stochastic Processes in Physics and Chemistry, North-Holland Personal Library, Amsterdam, 2007.

[13] M. E. J. Newman, Analysis of weighted networks, Physical Review E 70 (5) (2004) 056131. doi:10.1103/PhysRevE.70.056131.

[14] R. A. Horn, C. R. Johnson, Matrix Analysis, Cambridge University Press, New York, 2012.

[15] P. Bonacich, Power and centrality: A family of measures, American Journal of Sociology 92 (5) (1987) 1170–1182. doi:10.1086/228631.

[16] M. Newman, Networks, Oxford university press, Oxford, 2018.

[17] E. Abbe, Community detection and stochastic block models: Recent developments, Journal of Machine Learning Research 18 (177) (2018) 1–86.

[18] T. P. Peixoto, Bayesian Stochastic Blockmodeling, John Wiley & Sons, Ltd, Hoboken, 2019, Ch. 11, pp. 289–332. doi:10.1002/9781119483298.ch11.

[19] E. Côme, N. Jouvin, P. Latouche, C. Bouveyron, Hierarchical clustering with discrete latent variable models and the integrated classification likelihood, Advances in Data Analysis and Classification 15 (2021) 957–986. doi:10.1007/s11634-021-00440-z.

[20] U. Von Luxburg, A tutorial on spectral clustering, Statistics and Computing 17 (4) (2007) 395–416. doi:10.1007/s11222-007-9033-z.

[21] C. R. John, D. Watson, M. R. Barnes, C. Pitzalis, M. J. Lewis, Spectrum: fast density-aware spectral clustering for single and multi-omic data, Bioinformatics 36 (4) (2020) 1159–1166. doi:10.1093/bioinformatics/btz704.

[22] TCGA, The cancer genome atlas (2023). URL https://www.cancer.gov/tcga

[23] C. W. Brennan, R. G. Verhaak, A. McKenna, B. Campos, H. Noushmehr, S. R. Salama, et al., The somatic genomic landscape of glioblastoma, Cell 155 (2) (2013) 462–77. doi:10.1007/s00401-016-1545-1.

[24] TCGA, Comprehensive, integrative genomic analysis of diffuse lower-grade gliomas, New England Journal of Medicine 372 (2015) 2481–2498. doi:10.1056/NEJMoa1402121.

[25] TCGA, Comprehensive genomic characterization defines human glioblastoma genes and core pathways, Nature 455 (23) (2008) 1061–8. doi:10.1007/s00401-016-1545-1.

[26] D. N. Louis, H. Ohgaki, O. D. Wiestler, W. K. Cavenee, P. C. Burger, A. Jouvet, B. W. Scheithauer, P. Kleihues, The 2007 who classification of tumours of the central nervous system, Acta Neuropathologica 114 (2) (2007) 97–109. doi:10.1007/s00401-007-0243-4.

[27] J. Li, Y. Lu, R. Akbani, Z. Ju, P. L. Roebuck, W. Liu, J.-Y. Yang, B. M. Broom, R. G. W. Verhaak, D. W. Kane, C. Wakefield, J. N. Weinstein, G. B. Mills, H. Liang, TCPA: a resource for cancer functional proteomics data, Nature Methods 10 (11) (2013) 1046–1047. doi:10.1038/nmeth.2650.

[28] H. Liu, J. Lafferty, L. Wasserman, The nonparanormal: Semiparametric estimation of high dimensional undirected graphs, Journal of Machine Learning Research 10 (80) (2009) 2295–2328. URL http://jmlr.org/papers/v10/liu09a.html

[29] T. M. J. Fruchterman, E. M. Reingold, Graph drawing by force-directed placement, Software: Practice and Experience 21 (11) (1991) 1129–1164. doi:10.1002/spe.4380211102.

[30] L. Hubert, P. Arabie, Comparing partitions, Journal of Classification 2 (1) (1985) 193–218. doi:10.1007/BF01908075.

[31] L. Kaufman, P. Rousseeuw, Finding Groups in Data. An Introduction to Cluster Analysis, Wiley Inter-Science, New York, 1990.

[32] G. Reifenberger, V. P. Collins, Pathology and molecular genetics of astrocytic gliomas, Journal of Molecular Medicine 82 (2004) 656–670. doi:10.1007/s00109-004-0564-x.

[33] V. P. Andrade, M. Morrogh, L.-X. Qin, N. Olvera, D. Giri, S. Muhsen, R. A. Sakr, M. Schizas, C. K. Y. Ng, C. D. Arroyo, E. Brogi, A. Viale, M. Morrow, J. S. Reis-Filho, T. A. King, Gene expression profiling of lobular carcinoma in situ reveals candidate precursor genes for invasion, Molecular Oncology 9 (2015) 772–782. doi:10.1016/j.molonc.2014.12.005.

[34] J. Vallejo-Díaz, M. Chagoyen, M. Olazabal-Morán, A. González-García, A. C. Carrera, The opposing roles of pik3r1/p85α and pik3r2/p85,B in cancer, Trends in Cancer 5 (2019) 233–244. doi:10.1016/j.trecan.2019.02.009.

[35] Y. Liu, D. Wang, Z. Li, X. Li, M. Jin, N. Jia, X. Cui, G. Hu, T. Tang, Q. Yu, Pan-cancer analysis on the role of pik3r1 and pik3r2 in human tumors, Scientific Reports 12 (2022) 5924. doi:10.1038/s41598-022-09889-0.

[36] D. Chicco, A. Alameer, S. Rahmati, G. Jurman, Towards a potential pan-cancer prognostic signature for gene expression based on probesets and ensemble machine learning, BioData Mining 15 (2022) 28. doi:10.1186/s13040-022-00312-y.

[37] I. Wierstra, J. Alves, Foxm1, a typical proliferation-associated transcription factor, Biological Chemistry 388 (2007) 1257–1274. doi:10.1515/BC.2007.159.

[38] G.-B. Liao, X.-Z. Li, S. Zeng, C. Liu, S.-M. Yang, L. Yang, C.-J. Hu, J.-Y. Bai, Regulation of the master regulator foxm1 in cancer, Cell Communication and Signaling 16 (2018) 57. doi:10.1186/s12964-018-0266-6.

[39] S. Borhani, A. L. Gartel, Foxm1: a potential therapeutic target in human solid cancers, Expert Opinion on Therapeutic Targets 24 (2020) 205–217. doi:10.1080/14728222.2020.1727888.

[40] M. S. Chesnokov, S. Borhani, M. Halasi, Z. Arbieva, I. Khan, A. L. Gartel, Foxm1-akt positive regulation loop provides venetoclax resistance in aml, Frontiers in Oncology 11 (2021). doi:10.3389/fonc.2021.696532.

[41] X. Li, C. Wu, N. Chen, H. Gu, A. Yen, L. Cao, E. Wang, L. Wang, Pi3k/akt/mtor signaling pathway and targeted therapy for glioblastoma, Oncotarget 7 (22) (2016) 33440–33450. doi:10.18632/oncotarget.7961.

[42] E. Laurini, D. Marson, A. Fermeglia, S. Aulic, M. Fermeglia, S. Pricl, Role of rad51 and dna repair in cancer: A molecular perspective, Pharmacology & Therapeutics 208 (2020) 107492. doi:10.1016/j.pharmthera.2020.107492.

[43] H. O. King, T. Brend, H. L. Payne, A. Wright, T. A. Ward, K. Patel, T. Egnuni, L. F. Stead, A. Patel, H. Wurdak, S. C. Short, Rad51 is a selective dna repair target to radiosensitize glioma stem cells, Stem Cell Reports 8 (2017) 125–139. doi:10.1016/j.stemcr.2016.12.005.

[44] C. Morrison, E. Weterings, D. Mahadevan, A. Sanan, M. Weinand, B. Stea, Expression levels of rad51 inversely correlate with survival of glioblastoma patients, Cancers 13 (21) (2021). doi:10.3390/cancers13215358.

[45] S. Zona, L. Bella, M. J. Burton, G. N. de Moraes, E. W.-F. Lam, Foxm1: An emerging master regulator of dna damage response and genotoxic agent resistance, Biochimica et Biophysica Acta (BBA) - Gene Regulatory Mechanisms 1839 (2014) 1316–1322. doi:10.1016/j.bbagrm.2014.09.016.

[46] N. Zhang, X. Wu, L. Yang, F. Xiao, H. Zhang, A. Zhou, Z. Huang, S. Huang, Foxm1 inhibition sensitizes resistant glioblastoma cells to temozolomide by downregulating the expression of dna-repair gene rad51, Clinical Cancer Research 18 (2012) 5961–5971. doi:10.1158/1078-0432.CCR-12-0039.

[47] P. Tabnak, A. H. Bashkandi, M. Ebrahimnezhad, M. Soleimani, Fork-head box transcription factors (foxos and foxm1) in glioma: from molecular mechanisms to therapeutics, Cancer Cell International 23 (2023) 238. doi:10.1186/s12935-023-03090-7.

[48] S. Pal, D. Kozono, X. Yang, W. Fendler, W. Fitts, J. Ni, J. A. Alberta, J. Zhao, K. X. Liu, J. Bian, N. Truffaux, W. A. Weiss, A. C. Resnick, P. Bandopadhayay, K. L. Ligon, S. G. DuBois, S. Mueller, D. Chowdhury, D. A. Haas-Kogan, Dual hdac and pi3k inhibition abrogates nfkb- and foxm1-mediated dna damage response to radiosensitize pediatric high-grade gliomas, Cancer Research 78 (2018) 4007–4021. doi:10.1158/0008-5472.CAN-17-3691.

[49] V. C. Y. Mak, X. Li, L. Rao, Y. Zhou, S.-W. Tsao, L. W. T. Cheung, p85,B alters response to egfr inhibitor in ovarian cancer through p38 mapk-mediated regulation of dna repair, Neoplasia 23 (2021) 718–730. doi:10.1016/j.neo.2021.05.009.

[50] R. Coletti, M. L. Mendonça, S. Vinga, M. B. Lopes, Inferring diagnostic and prognostic gene expression signatures across WHO glioma classifications: A network-based approach (2023). doi:10.48550/arXiv.2305.12207.

[51] T. Schlitt, A. Brazma, Current approaches to gene regulatory network modelling, BMC Bioinformatics 8 (S6) (2007) S9. doi:10.1186/1471-2105-8-S6-S9.

[52] K. Bhattacharjee, M. Nath, Y. Choudhury, Fatty acid synthesis and cancer: Aberrant expression of the acaca and acacb genes increases the risk for cancer, Meta Gene 26 (2020) 100798. doi:10.1016/j.mgene.2020.100798.

[53] H. Zhang, S. Liu, Z. Cai, W. Dong, J. Ye, Z. Cai, Z. Han, Y. Liang, Y. Zhuo, Y. Luo, X. Zhu, Y. Deng, Y. Zhang, R. Liu, Y. Feng, J. Lai, R. Zhou, H. Tan, W. Zhong, Down-regulation of acaca suppresses the malignant progression of prostate cancer through inhibiting mitochondrial potential, Journal of Cancer 12 (1) (2021) 232–243. doi:10.7150/jca.49560.

[54] R. Malla, C. R. Ashby, N. K. Narayanan, B. Narayanan, J. S. Faridi, A. K. Tiwari, Proline-rich AKT substrate of 40-kDa (PRAS40) in the pathophysiology of cancer, Biochemical and Biophysical Research Communications 463 (3) (2015) 161–166. doi:10.1016/j.bbrc.2015.05.041.

[55] Y. Zhao, Q. Zhao, P. J. Kaboli, J. Shen, M. Li, X. Wu, J. Yin, H. Zhang, Y. Wu, L. Lin, L. Zhang, L. Wan, Q. Wen, X. Li, C. H. Cho, T. Yi, J. Li, Z. Xiao, m1A Regulated Genes Modulate PI3K/AKT/mTOR and ErbB Pathways in Gastrointestinal Cancer, Translational Oncology 12 (10) (2019) 1323–1333. doi:10.1016/j.tranon.2019.06.007.

[56] K. M. Turner, Y. Sun, P. Ji, K. J. Granberg, B. Bernard, L. Hu, D. E. Cogdell, X. Zhou, O. Yli-Harja, M. Nykter, I. Shmulevich, W. K. A. Yung, G. N. Fuller, W. Zhang, Genomically amplified akt3 activates dna repair pathway and promotes glioma progression, Proceedings of the National Academy of Sciences 112 (11) (2015) 3421–3426. doi:10.1073/pnas.1414573112.

[57] A. Joy, M. Kapoor, J. Georges, L. Butler, Y. Chang, C. Li, A. Crouch, I. Smirnov, M. Nakada, J. Hepler, M. Marty, B. G. Feuerstein, The role of akt isoforms in glioblastoma: Akt3 delays tumor progression, Journal of Neuro-Oncology 130 (1) (2016) 43–52. doi:10.1007/s11060-016-2220-z.

[58] J. M. Abdullah, F. Ahmad, K. A. K. Ahmad, M. M. Ghazali, H. Jaafar, A. Ideris, A. M. Ali, A. R. Omar, K. Yusoff, M. A. M. Lila, F. Othman, Molecular genetic analysis of BAX and cyclin D1 genes in patients with malignant glioma, Neurological Research 29 (3) (2007) 239–242. doi:10.1179/016164107X158965.

[59] J. Huang, J. Gao, X. Lv, G. Li, D. Hao, X. Yao, L. Zhou, D. Liu, R. Wang, Target gene therapy of glioma: overexpression of bax gene under the control of both tissue-specific promoter and hypoxia-inducible element, Acta Biochimica et Biophysica Sinica 42 (4) (2010) 274–280. doi:10.1093/abbs/gmq016.

[60] C. L. Appin, D. J. Brat, Molecular genetics of gliomas, The Cancer Journal 20 (1) (2014) 66–72. doi:10.1097/PPO.0000000000000020.

[61] H. G. Vuong, A. M. A. Altibi, U. N. P. Duong, H. T. T. Ngo, T. Q. Pham, K.-M. Fung, L. Hassell, Braf mutation is associated with an improved survival in glioma—a systematic review and meta-analysis, Molecular Neurobiology (2017). doi:10.1007/s12035-017-0599-y.

[62] S. Quiros, W. P. Roos, B. Kaina, Rad51 and brca2 - new molecular targets for sensitizing glioma cells to alkylating anticancer drugs, PLoS ONE 6 (11) (2011) e27183. doi:10.1371/journal.pone.0027183.

[63] S. E. Meimand, A. Pour-Rashidi, M. M. Shahrbabak, E. Mohammadi, F. E. Meimand, N. Rezaei, The prognostication potential of brca genes expression in gliomas: A genetic survival analysis study, World Neurosurgery 157 (2022) e123–e128. doi:10.1016/j.wneu.2021.09.107.

[64] S. Huang, J. Xiao, J. Wu, J. Liu, X. Feng, C. Yang, D. Xiang, S. Luo, Tizoxanide Promotes Apoptosis in Glioblastoma by Inhibiting CDK1 Activity, Frontiers in Pharmacology 13 (2022) 895573. doi:10.3389/fphar.2022.895573.

[65] L. Bo, B. Wei, C. Li, Z. Wang, Z. Gao, Z. Miao, Identification of potential key genes associated with glioblastoma based on the gene expression profile, Oncology Letters 14 (2) (2017) 2045–2052. doi:10.3892/ol.2017.6460.

[66] P. Apostolou, I. Papasotiriou, Current perspectives on chek2 mutations in breast cancer, Breast Cancer: Targets and Therapy 9 (2017) 331–335. doi:10.2147/BCTT.S111394.

[67] M. Simon, M. Ludwig, R. Fimmers, R. Mahlberg, A. Müller-Erkwoh, G. Köster, J. Schramm, Variant of the chek2 gene as a prognostic marker in glioblastoma multiforme, Neurosurgery 59 (5) (2006) 1078–1085. doi:10.1227/01.NEU.0000245590.08463.5B.

[68] Y.-L. Cha, P.-D. Li, L.-J. Yuan, M.-Y. Zhang, Y.-J. Zhang, H.-L. Rao, H.-Z. Zhang, X. F. S. Zheng, H.-Y. Wang, Eif4ebp1 overexpression is associated with poor survival and disease progression in patients with hepatocellular carcinoma, PLOS ONE 10 (2) (2015) e0117493. doi:10.1371/journal.pone.0117493.

[69] L. Hauffe, D. Picard, J. Musa, M. Remke, T. G. P. Grünewald, B. Rotblat, G. Reifenberger, G. Leprivier, Eukaryotic translation initiation factor 4e binding protein 1 (eif4ebp1) expression in glioblastoma is driven by ets1- and mybl2-dependent transcriptional activation, Cell Death Discovery 8 (1) (2022) 91. doi:10.1038/s41420-022-00883-z.

[70] J. Song, D. Zhao, G. Sun, J. Yang, Z. Lv, B. Jiao, PTPRM methylation induced by FN1 promotes the development of glioblastoma by activating STAT3 signalling, Pharmaceutical Biology 59 (1) (2021) 902–909. doi:10.1080/13880209.2021.1944220.

[71] S.-X. Cheng, Y. Tu, S. Zhang, FoxM1 Promotes Glioma Cells Progression by Up-Regulating Anxa1 Expression, PLoS ONE 8 (8) (2013) e72376. doi:10.1371/journal.pone.0072376.

[72] M. Liu, B. Dai, S.-H. Kang, K. Ban, F.-J. Huang, F. F. Lang, K. D. Aldape, T.-x. Xie, C. E. Pelloski, K. Xie, R. Sawaya, S. Huang, FoxM1B Is Overexpressed in Human Glioblastomas and Critically Regulates the Tumorigenicity of Glioma Cells, Cancer Research 66 (7) (2006) 3593–3602. doi:10.1158/0008-5472.CAN-05-2912.

[73] L. E. Huang, A. L. Cohen, H. Colman, R. L. Jensen, D. W. Fults, W. T. Couldwell, IGFBP2 expression predicts IDH-mutant glioma patient survival, Oncotarget 8 (1) (2017) 191–202. doi:10.18632/oncotarget.13329.

[74] D. Hsieh, A. Hsieh, B. Stea, R. Ellsworth, IGFBP2 promotes glioma tumor stem cell expansion and survival, Biochemical and Biophysical Research Communications 397 (2) (2010) 367–372. doi:10.1016/j.bbrc.2010.05.145.

[75] Z. Duzgun, Z. Eroglu, C. Biray Avci, Role of mTOR in glioblastoma, Gene 575 (2) (2016) 187–190. doi:10.1016/j.gene.2015.08.060.

[76] D. Akhavan, T. F. Cloughesy, P. S. Mischel, mtor signaling in glioblastoma: lessons learned from bench to bedside, Neuro-Oncology 12 (8) (2010) 882–889. doi:10.1093/neuonc/noq052.

[77] L. Jiang, J. Wu, Q. Chen, X. Hu, W. Li, G. Hu, Notch1 expression is upregulated in glioma and is associated with tumor progression, Journal of Clinical Neuroscience 18 (3) (2011) 387–390. doi:10.1016/j.jocn.2010.07.131.

[78] J. Li, Y. Cui, G. Gao, Z. Zhao, H. Zhang, X. Wang, Notch1 is an independent prognostic factor for patients with glioma, Journal of Surgical Oncology 103 (8) (2011) 813–817. doi:10.1002/jso.21851.

[79] G. Musumeci, A. Castorina, G. Magro, V. Cardile, S. Castorina, D. Ribatti, Enhanced expression of cd31/platelet endothelial cell adhesion molecule 1 (pecam1) correlates with hypoxia inducible factor-1 alpha (hif-1α) in human glioblastoma multiforme, Experimental Cell Research 339 (2) (2015) 407–416. doi:10.1016/j.yexcr.2015.09.007.

[80] C. K. Cheng, Q. Fan, W. A. Weiss, Pi3k signaling in glioma—animal models and therapeutic challenges, Brain Pathology 19 (1) (2009) 112–120. doi:10.1111/j.1750-3639.2008.00233.x.

[81] A. Zajaç, et al., Involvement of pi3k pathway in glioma cell resistance to temozolomide treatment, International Journal of Molecular Sciences 22 (10) (2021) 5155. doi:10.3390/ijms22105155.

[82] J. Langhans, L. Schneele, N. Trenkler, H. Von Bandemer, L. Nonnenmacher, G. Karpel-Massler, M. D. Siegelin, S. Zhou, M.-E. Halatsch, K.-M. Debatin, M.-A. Westhoff, The effects of pi3k-mediated signalling on glioblastoma cell behaviour, Oncogenesis 6 (11) (2017) 398. doi:10.1038/s41389-017-0004-8.

[83] J. Mathivanan, K. Rohini, M. L. Gope, B. Anandh, R. Gope, Altered structure and deregulated expression of the tumor suppressor gene retinoblastoma (rb1) in human brain tumors, Molecular and Cellular Biochemistry 302 (1-2) (2007) 67–77. doi:10.1007/s11010-007-9428-3.

[84] Y. Kim, J. Lachuer, M. Mittelbronn, W. Paulus, B. Brokinkel, K. Keyvani, U. Sure, K. Wrede, S. Nobusawa, Y. Nakazato, Y. Tanaka, A. Vital, L. Mariani, H. Ohgaki, Alterations in the RB1 Pathway in Low-grade Diffuse Gliomas Lacking Common Genetic Alterations, Brain Pathology 21 (6) (2011) 645–651. doi:10.1111/j.1750-3639.2011.00492.x.

[85] M. Nakamura, Y. Yonekawa, P. Kleihues, H. Ohgaki, Promoter Hyper-methylation of the RB1 Gene in Glioblastomas, Laboratory Investigation 81 (1) (2001) 77–82. doi:10.1038/labinvest.3780213.

[86] Y. Shirakawa, T. Hide, M. Yamaoka, Y. Ito, N. Ito, K. Ohta, N. Shinojima, A. Mukasa, H. Saito, H. Jono, Ribosomal protein S6 promotes stem-like characters in glioma cells, Cancer Science 111 (6) (2020) 2041–2051. doi:10.1111/cas.14399.

[87] Y. Shirakawa, K. Ohta, S. Miyake, A. Kanemaru, A. Kuwano, K. Yonemaru, S. Uchino, M. Yamaoka, Y. Ito, N. Ito, T. Hide, N. Shinojima, A. Mukasa, H. Saito, H. Jono, Glioma Cells Acquire Stem-like Characters by Extrinsic Ribosome Stimuli, Cells 10 (11) (2021) 2970. doi:10.3390/cells10112970.

[88] T. Hide, I. Shibahara, M. Inukai, R. Shigeeda, Y. Shirakawa, H. Jono, N. Shinojima, A. Mukasa, T. Kumabe, Ribosomal proteins induce stem cell-like characteristics in glioma cells as an “extra-ribosomal function”, Brain Tumor Pathology 39 (2) (2022) 51–56. doi:10.1007/s10014-022-00434-5.

[89] F. Seker, et al., Identification of SERPINE1 as a Regulator of Glioblastoma Cell Dispersal with Transcriptome Profiling, Cancers 11 (11) (2019) 1651. doi:10.3390/cancers11111651.

[90] X. Huang, F. Zhang, D. He, X. Ji, J. Gao, W. Liu, Y. Wang, Q. Liu, T. Xin, Immune-Related Gene SERPINE1 Is a Novel Biomarker for Diffuse Lower-Grade Gliomas via Large-Scale Analysis, Frontiers in Oncology 11 (2021) 646060. doi:10.3389/fonc.2021.646060.

[91] X. Tan, L. Zou, J. Qin, D. Xia, Y. Zhou, G. Jin, Z. Jiang, H. Li, SQSTM1/p62 is involved in docosahexaenoic acid–induced cellular autophagy in glioblastoma cell lines, In Vitro Cellular & Developmental Biology - Animal 55 (9) (2019) 703–712. doi:10.1007/s11626-019-00387-8.

[92] A. Puissant, N. Fenouille, P. Auberger, When autophagy meets cancer through p62/sqstm1, Am J Cancer Res 2 (4) (2012) 397–413.

[93] V. Schumacher, STK11 genotyping and cancer risk in Peutz-Jeghers syndrome, Journal of Medical Genetics 42 (5) (2005) 428–435. doi:10.1136/jmg.2004.026294.

[94] V. Launonen, Mutations in the human LKB1/STK11 gene, Human Mutation 26 (4) (2005) 291–297. doi:10.1002/humu.20222.

[95] H. Wang, W. Li, G. Wang, S. Zhang, L. Bie, Overexpression of STMN1 is associated with the prognosis of meningioma patients, Neuroscience Letters 654 (2017) 1–5. doi:10.1016/j.neulet.2017.06.020.

[96] B. Belletti, G. Baldassarre, Stathmin: a protein with many tasks. new biomarker and potential target in cancer, Expert Opinion on Therapeutic Targets 15 (11) (2011) 1249–1266. doi:10.1517/14728222.2011.620951.

[97] G. Moncayo, M. Grzmil, T. Smirnova, P. Zmarz, R. M. Huber, D. Hynx, H. Kohler, Y. Wang, H.-R. Hotz, N. E. Hynes, G. Keller, S. Frank, A. Merlo, B. A. Hemmings, Syk inhibition blocks proliferation and migration of glioma cells and modifies the tumor microenvironment, Neuro-Oncology 20 (5) (2018) 621–631. doi:10.1093/neuonc/noy008.

[98] Q. Zhou, M. Wei, W. Shen, S. Huang, J. Fan, H. Huang, SYK Is Associated With Malignant Phenotype and Immune Check-points in Diffuse Glioma, Frontiers in Genetics 13 (2022) 899883. doi:10.3389/fgene.2022.899883.

